## Supplementary material for "Understanding the mechanism of facilitation in hoverfly TSDNs": S1 Appendix

### Supporting information

##### S1 Appendix. Implementation details of the STMD model.

##### Early visual processing

Similarly to Wiederman et al. [1], our ESTMD model only used the green channel of the RGB image input – this is an approximation of the green spectral sensitivity of the movement detection mechanisms in hoverflies [2]. The tiny lenses of compound eyes cause diffraction of light [3], which results in reduced spatial acuity, i.e., high optical blur. Following Wiederman et al. [1], we modelled this optical blur by convolving a circularly symmetric 2D Gaussian kernel with the stimulus image,

$$G(x,y) = \frac{1}{2\pi\sigma^2} \exp\left(-\frac{(x^2+y^2)}{2\sigma^2}\right) \quad (1)$$

where $\sigma$ is the standard deviation. In order to emulate the hoverfly optics [4], we calculated $\sigma$ through the full-width at half maximum (FWHM) of the fly optical angular sensitivity, which is 1.4° [3]. Therefore, $\sigma$ can be calculated in pixels as:

$$\sigma = \frac{1.4}{2\sqrt{2\ln 2}} \times \text{pixels/degree} \approx 16.7 \text{ pixels} \quad (2)$$

where pixels/degree = 2560/155 is calculated by dividing the spatial resolution of the visual stimulus by the subtended viewing angle of the visual field of the hoverfly eye, based on an inter-ommatidial angle of 1° [4]. This consideration is also applied next for the spatial downsampling of the blurred image, where each pixel represents a degree of the visual field. The kernel size of the discrete Gaussian approximation is determined as $2\lceil 3\sigma \rceil - 1 = 99$ pixels, as opposed to the kernel size determination used by Bagheri et al. [5], which gives $2\lceil \sigma \rceil = 34$ pixels. Our choice of kernel size results in a discrete kernel that includes the $99^{th}$ percentile of the Gaussian, while the odd size mitigates aliasing.

Next, the Lamina Monopolar Cells (LMCs) remove redundant information by using neuronal adaptation (temporal high-pass filtering) and centre-surround antagonism (spatial high-pass filtering). Wiederman et al. [1] modelled this using a Lipetz transform. Since the temporal filtering qualities of both the photoreceptors and LMC can be combined to give a temporal band-pass filter [5], for simplicity, we used such a filter instead. This filter was approximated and simulated using a discrete transfer function of a difference of log-normals (DLN) as per the method described by Halupka et al. [6]:

$$y[n] = \sum_{i=1}^{8} \alpha_i x[n+(i-9)] - \sum_{i=1}^{8} \beta_i y[n+(i-9)] \quad (3)$$

where $x[n]$ is the unfiltered signal and $y[n]$ the filtered output at a discrete timestep, $n$, and $\alpha_i$ and $\beta_i$ are the numerator and denominator coefficients given in Table A.

Again, similar to [5], we applied centre-surround antagonism in the LMC by convolving the image with a zero-mean high-pass filter kernel $H$:

$$H = \frac{1}{9}\begin{bmatrix} -1 & -1 & -1 \\ -1 & 8 & -1 \\ -1 & -1 & -1 \end{bmatrix}$$

**Table A.** Coefficients of the discrete DLN function shown in Eq 3.

| Numerator coefficients | | Denominator coefficients | |
|---|---|---|---|
| $\alpha_1$ | $-0.15240$ | $\beta_1$ | 0.06510 |
| $\alpha_2$ | 0.17890 | $\beta_2$ | $-0.54180$ |
| $\alpha_3$ | $-0.05700$ | $\beta_3$ | 2.14480 |
| $\alpha_4$ | 0.04300 | $\beta_4$ | $-5.30600$ |
| $\alpha_5$ | $-0.01600$ | $\beta_5$ | 9.00040 |
| $\alpha_6$ | 0.00440 | $\beta_6$ | $-10.71100$ |
| $\alpha_7$ | $-0.00076$ | $\beta_7$ | 8.68500 |
| $\alpha_8$ | 0.00006 | $\beta_8$ | $-4.33300$ |

##### Target matched filtering

Similar to [1, 5], the output from the LMC was first half-wave rectified and then separated into ON (brightness increments) and OFF (brightness decrements) contrast polarity channels:

$$\mathrm{I}_{\mathrm{ON}}(y) = \begin{cases} y & \text{if } y > 0 \\ 0 & otherwise \end{cases} \tag{4}$$

$$\mathrm{I}_{\mathrm{OFF}}(y) = \begin{cases} -y & \text{if } y < 0 \\ 0 & otherwise \end{cases} \tag{5}$$

where $y$ represents the output from the LMC at a discrete timestep and $I$ is the independent channel half-wave rectified output, i.e., ON or OFF channel. Wiederman et al. [1] was inspired by the rectifying transient cells (RTCs) found in the blowfly medulla [7]. This fast-adaptive mechanism, which 'adapts' the independent ON and OFF channels, was also implemented in [5] and this work. Depending on whether the independent channel signal is increasing or decreasing, the time constant of the nonlinear filter (NLF) used to implement this process switches [1]. The time constant is 'fast' ($\tau_{\text{fast}} = 10\,\text{ms}$) for an increasing signal or 'slow' ($\tau_{\text{slow}} = 100\,\text{ms}$) for a decreasing signal. Subsequently, for each polarity channel, at each discrete timestep, the channel NLF value is subtracted from the half-wave rectified channel signal. Through this subtraction, responses to the contrasting edges of visual features are enhanced, whilst lower-contrast 'textural details' are rejected. The NLF is computed through a temporal low-pass filter,

$$\mathrm{NLF}[n] = (1 - \alpha[n]) \cdot I[n] + \alpha[n] \cdot \mathrm{NLF}[n-1] \tag{6}$$

where $I[n]$ represents the independent channel signal at a certain discrete timestep $n$. The smoothing factor, $\alpha$ of the NLF is selected using the fast-adaptive mechanism described above:

$$\alpha[n] = \begin{cases} \exp\left(-T_s/\tau_{\text{fast}}\right) & \text{if } I[n] - I[n-1] > 0 \\ \exp\left(-T_s/\tau_{\text{slow}}\right) & \text{if } I[n] - I[n-1] \le 0 \end{cases} \tag{7}$$

where $T_s$ is the sampling period ($T_s = 1\,\text{ms}$). While Wiederman et al. [1] used $\tau_{\text{fast}} = 1\,\text{ms}$, $\tau_{\text{slow}} = 100\,\text{ms}$ and Bagheri et al. [5] used $\tau_{\text{fast}} = 3\,\text{ms}$, $\tau_{\text{slow}} = 70\,\text{ms}$, we used $\tau_{\text{fast}} = 10\,\text{ms}$, $\tau_{\text{slow}} = 100\,\text{ms}$ as we found that these parameters led to ESTMD spatio-temporal tuning characteristics that better matched hoverfly physiology [8]. Similar to both [1, 5], the independently adapted ON and OFF channels were then convolved with a further centre-surround antagonistic kernel $H_{\text{inh}}$. This process represents the spatial lateral inhibitory effect within polarity channels seen in blowfly

RTCs [9]. Target size tuning is achieved by varying the gain, $\gamma$ and spatial extent of the centre-surround antagonism described by the $H_{\text{inh}}$ kernel:

$$H_{\text{inh}} = \gamma \cdot \begin{bmatrix} -1 & -1 & -1 & -1 & -1 \\ -1 & 0 & 0 & 0 & -1 \\ -1 & 0 & 2 & 0 & -1 \\ -1 & 0 & 0 & 0 & -1 \\ -1 & -1 & -1 & -1 & -1 \end{bmatrix} \tag{8}$$

While Wiederman et al. [1] accomplished spatial lateral antagonism by subtracting a temporally low-pass filtered signal of the surrounding pixels from the independent channel signal, $I[n]$, the spatial extent of this antagonism, or weighting of the extent, was not addressed. Instead, Bagheri et al. [5] presented a centre-surround kernel M (Figs 4, 9 of [5]) that led to their ESTMD model having a band-pass region responsive to targets similar to the ones used in this work. Eq 8 is the same kernel as kernel M. The laterally inhibited independent ON and OFF channels were then clamped above zero to ensure the channel signals were non-negative. Similar to [1, 5], the local OFF channel outputs were delayed before correlating with undelayed local ON channel outputs to form a 'matched filter'. The OFF channels were delayed to enhance selectivity for dark targets. The matched filter formulation in the discrete-time domain is ($\tau = 25\,\text{ms}$, $T_s = 1\,\text{ms}$):

$$\text{ESTMD Output}[n] = ON[n] \cdot \text{D}(OFF[n]) \tag{9}$$

The OFF channel is temporally low-pass filtered – the delay operator, D was obtained using a temporal low-pass filter, similar to the one described in Eq 6.
